# Magnetic Levitation Derived Metabolomic Fingerprinting Enables Exploratory Discrimination of Breast Cancer Subtypes

**DOI:** 10.64898/2026.05.23.727414

**Authors:** Samantha Velazquez, Colin C. Anderson, Julie A. Reisz, Ali Akbar Ashkarran

## Abstract

Breast cancer (BC) consists of heterogeneous molecular subtypes with distinct biological behavior and therapeutic response, including triple negative breast cancer (TNBC), human epidermal growth factor receptor 2 positive (HER2+), and Luminal A tumors. Current subtype classification primarily relies on tissue biopsy and molecular pathology, highlighting the need for complementary non invasive analytical approaches capable of capturing systemic biochemical differences among BC subtypes. In this exploratory study, we investigate whether magnetic levitation (MagLev) derived plasma patterning combined with untargeted metabolomics analysis can distinguish BC subtypes based on subtype specific metabolomic fingerprints. Representative plasma samples from TNBC, HER2+, and Luminal A patients were levitated in a standard MagLev system containing superparamagnetic iron oxide nanoparticles (SPIONs), generating visibly distinct levitation patterns throughout the levitation process. Individual levitated plasma layers were subsequently extracted and subjected to untargeted metabolomics analysis. Multivariate analysis demonstrated clear subtype dependent metabolic separation among the three BC subtypes. Heatmap clustering, PLS DA analysis, and variable importance profiling identified distinct metabolic signatures involving glycolysis, pentose phosphate pathway metabolism, TCA cycle activity, amino acid metabolism, and lipid remodeling. Elevated levels of phosphoglycerate, pyruvate, ribose 5 phosphate, glutamate, and 2 oxoglutarate suggested enhanced proliferative and biosynthetic metabolism, while enrichment of long chain fatty acids in Luminal A samples indicated subtype specific lipid metabolic remodeling. These findings demonstrate the feasibility of combining MagLev derived plasma organization with metabolomics analysis to generate disease specific metabolomic fingerprints and establish a foundation for future large scale validation studies.

## Introduction

Breast cancer remains one of the most common malignancies worldwide and exhibits substantial molecular and clinical heterogeneity.[1-3] Among the major molecular subtypes, triple negative breast cancer (TNBC), human epidermal growth factor receptor 2 positive (HER2+), and Luminal A tumors display distinct biological behavior, prognosis, and therapeutic response. TNBC lacks expression of estrogen receptor (ER), progesterone receptor (PR), and HER2, resulting in aggressive progression and limited targeted treatment options.[4-8] HER2+ tumors are characterized by HER2 overexpression and altered proliferative signaling, while Luminal A tumors generally display hormone receptor positivity and less aggressive clinical behavior. Despite advances in molecular pathology and receptor-based classification, current diagnostic workflows still rely heavily on tissue biopsy and invasive procedures.[9-11]

Emerging evidence suggests that cancer progression induces systemic biochemical and metabolic alterations that can potentially be captured through analysis of biological fluids such as plasma.[12, 13] Metabolomics analysis has therefore emerged as a promising strategy for identifying disease specific metabolic signatures and pathway alterations associated with tumor biology. Several studies have demonstrated that cancer cells exhibit altered glycolysis, pentose phosphate pathway activity, TCA cycle remodeling, amino acid metabolism, and lipid metabolism to support rapid proliferation and biosynthesis.[14-16] However, direct plasma metabolomics analysis often faces challenges related to the complexity and dynamic range of plasma biomolecules.[17, 18]

Magnetic levitation (MagLev) has recently emerged as a powerful platform for density based manipulation, separation, and characterization of diamagnetic materials, including complex biological systems and biomolecules.[19-28] Over the past decade, MagLev has attracted increasing attention in biological and biomedical research for applications ranging from cell sorting and tissue engineering to biomaterial assembly and regenerative medicine.[29-35] Recent studies have demonstrated the potential of MagLev as a label free analytical platform for disease detection and biological phenotyping. [23, 26, 28, 36-42] For example, MagLev systems have been used for differentiation of healthy and diseased blood cells, identification of circulating tumor cells, and detection of intracellular lipid accumulation based on subtle density and biophysical differences among cells and biomolecular systems.[43-46] We have previously shown that MagLev platforms can be integrated with downstream molecular analysis workflows for biological detection applications, highlighting their potential as versatile tools for disease related biomolecular characterization.

MagLev systems provide a unique platform in which complex biological samples respond to externally applied magnetic field gradient according to their density, magnetic susceptibility, and biomolecular composition.[47, 48] As a result of these interactions, biomolecules and biological components become spatially organized into distinct and dynamically evolving patterns or layers within the MagLev system.[49] We therefore hypothesized that disease specific plasma samples may generate unique spatially resolved biomolecular organizations that can be further analyzed to identify disease dependent molecular fingerprints.[50, 51] We have previously demonstrated that plasma samples introduced into MagLev systems generate dynamic levitation patterns consisting of distinct ellipsoidal bands or layers that evolve over time. These levitated layers may contain subtype dependent biomolecular information that can be exploited for disease discrimination.[49-51] In this exploratory proof-of-concept study, we investigate whether MagLev derived plasma patterning combined with untargeted metabolomics analysis can distinguish among three major BC subtypes including TNBC, HER2+, and Luminal A, which represent biologically distinct forms BC. By integrating physical plasma organization within a magnetic field with downstream metabolomics analysis, we aim to establish the feasibility of generating subtype specific metabolomic fingerprints for BC discrimination.

## Experimental details

### Materials

SPIONs (30 mg/mL) functionalized with ferumoxytol were purchased from Feraheme (www.feraheme.com) and diluted with phosphate buffered saline (PBS, 1X) solution to desired concentrations. Fluorescent polyethylene microparticles with known densities were obtained from Cospheric (www.cospheric.com). Three major subtypes (n = 1 per subtype) of BC human plasma samples were ordered from Innovative Research and were normalized to a concentration of 35 mg/ml (PBS, 1X) for all experiments.

### MagLev platform

The schematic representation and photograph of the standard MagLev platform used in this study is shown in **Figures 1a** and **2c**. The system consisted of two N42 grade neodymium (NdFeB) cubic magnets (25.4 mm × 25.4 mm × 50.8 mm ordered from K&J magnetics) positioned with like poles facing each other in an anti-Helmholtz configuration. The magnets were separated by a fixed distance of 2.5 cm. It is also noteworthy that this separation distance was optimized based on our previous studies to achieve a linear relationship between sample density and levitation height.[49] **Figure 2b** shows the linear relationship between sample density and levitation height using standard density glass beads at different SPION concentrations. Disposable plastic cuvettes trimmed to 25 mm were used as levitation container. Similar to previously reported MagLev systems, the platform generates a magnetic field gradient between two identical coaxial permanent magnets with face-to-face identical poles separated by a distance d.[52] In the anti-Helmholtz configuration, the magnetic field is primarily directed along the z axis, while the magnetic field components in the XY plane cancel each other, resulting in magnetic field lines that appear to repel each other (**Figure 2c**). This configuration generates a strong magnetic field gradient that drives objects toward regions of lower magnetic field strength and higher magnetic field gradients, enabling density-based levitation and separation. **Figure 2d** shows the levitation profiles of mixed approximately 40 to 50 µm fluorescent polyethylene microspheres with different densities over time.

**Figure 1:**
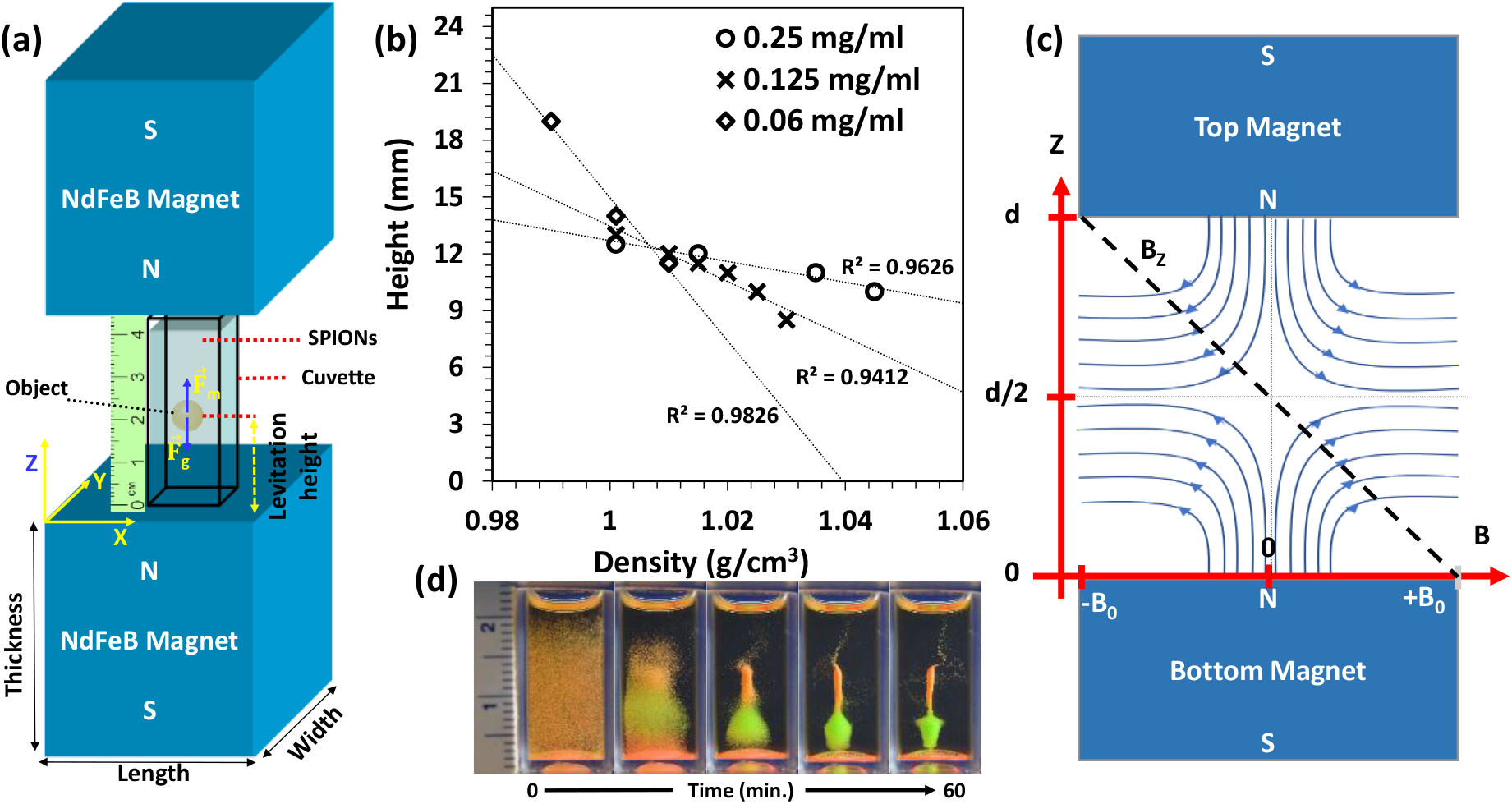
(a) Schematic showing the configuration of standard MagLev system, (b) calibration curves of the MagLev system showing the linear relation between density and levitation height at different concentrations of SPIONs, (c) the pattern of the magnetic field lines between the magnets, and (d) levitation profiles of the mixed polyethylene fluorescent microspheres (∼ 40-50 µm) and densities of 1.011 g/cm^3^ (orange) and 1.033 g/cm^3^ (green) over the time.

**Figure 2:**
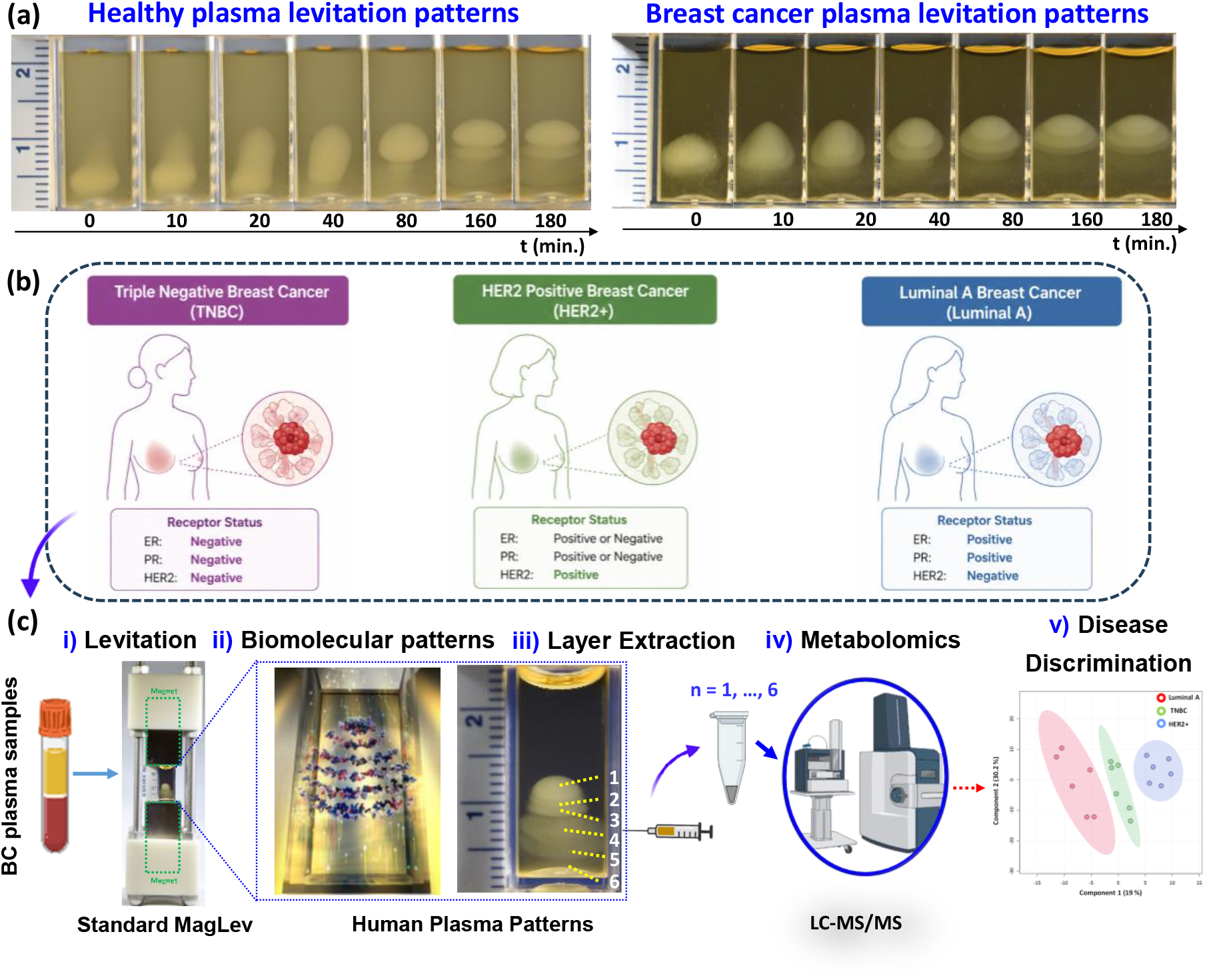
(a) Representative images showing the levitation profiles of healthy and BC human plasmas, (b) scheme showing the three different types of BC used in this study as a proof-of-concept model, and (c) overall workflow of the study for BC subtypes discrimination using MagLev system and metabolomics analysis.

### Levitation of the BC plasma samples

Plasma samples were levitated in a standard MagLev system using 0.06 mg/mL SPIONs, an optimized concentration based on our previous studies.[49-51] For all experiments (minimum n = 3 replicates), 100 μL of normalized plasma sample was injected into the MagLev solution and monitored throughout the levitation process for up to 3 hours. To minimize variability among plasma samples, all plasma protein concentrations were normalized to 35 mg/mL using a bicinchoninic acid (BCA) assay prior to levitation. Following formation of the levitated plasma patterns, individual ellipsoidal layers were sequentially extracted from top to bottom layer by layer using a plastic cuvette with several holes in the sidewall and a 0.5 ml insulin syringe under controlled negative pressure and stored for metabolomics analysis.

### Untargeted metabolomics analysis

Following magnetic levitation and extraction of the levitated plasma layers, metabolites were extracted using a cold solvent mixture consisting of methanol, acetonitrile, and water (5:3:2, v/v/v) added at a 1:25 sample dilution ratio. Samples were vortexed vigorously for 30 min at 4 °C to ensure efficient metabolite extraction and protein precipitation, followed by centrifugation at 18,213 rcf for 10 min at 4 °C. The resulting supernatants were transferred to LC MS vials for analysis. Metabolite separation and detection were carried out using 10 μL injection volumes under standardized LC MS conditions. Raw mass spectrometry files were converted to mzXML format using RawConverter software prior to downstream processing. Metabolite assignment and peak integration were performed using Maven in conjunction with the KEGG metabolomics database and an in house metabolite standard library. Chromatographic alignment, peak extraction, normalization, and multivariate statisin-housealyses were conducted using MetaboAnalyst workflows. Principal component analysis (PCA), partial least squares discriminant analysis (PLS DA), heatmap clustering, volcano plots, and variable importance projection (VIP) analyses were used to evaluate subtype dependent metabolomic differences among the BC plasma samples.

## Results and discussion

Figure 2. illustrates the overall workflow of the study. Three major and biologically heterogeneous BC subtypes including TNBC, HER2+, and Luminal A were selected as a proof-of-concept and representative models in this study. Individual levitated plasma patterns and the corresponding layers were subsequently extracted from top to bottom and subjected to untargeted LC MS metabolomics analysis to identify subtype specific metabolomic fingerprints associated with each BC subtype. Our previous findings demonstrated that human plasma forms distinct and highly reproducible ellipsoidal biomolecular patterns when levitated in a MagLev system containing SPIONs (**Figures 2–3**).[49-51] Pattern formation initiates within minutes after plasma injection and dynamically evolves over approximately 3 hours, ultimately generating stable and spatially resolved multilayer structures. Importantly, these characteristic levitation patterns are observed only for complex plasma systems and not for individual proteins or simple protein mixtures, suggesting that the patterns arise from collective physicochemical interactions among plasma biomolecules under externally applied magnetic field gradients.[49] Moreover, pattern formation occurs exclusively within the MagLev system and in the presence of magnetic fields, further supporting the role of magnetic field driven plasma organization in generating disease related biomolecular fingerprints.

Representative levitation images of normalized plasma samples from three BC subtypes in 0.06 mg/mL SPIONs within the standard MagLev system (**Figure 3**) revealed visibly distinct levitation patterns throughout the levitation process (0–180 min), with subtype specific differences observed not only in the final plasma organization but also during the dynamic evolution of the patterns over time, suggesting: **(i)** plasma pattern formation is not limited to a single disease condition and **(ii)** the feasibility of MagLev-based BC subtype discrimination. Although preliminary and limited to representative samples, these findings support the feasibility of the proposed larger scale investigation.

**Figure 3:**
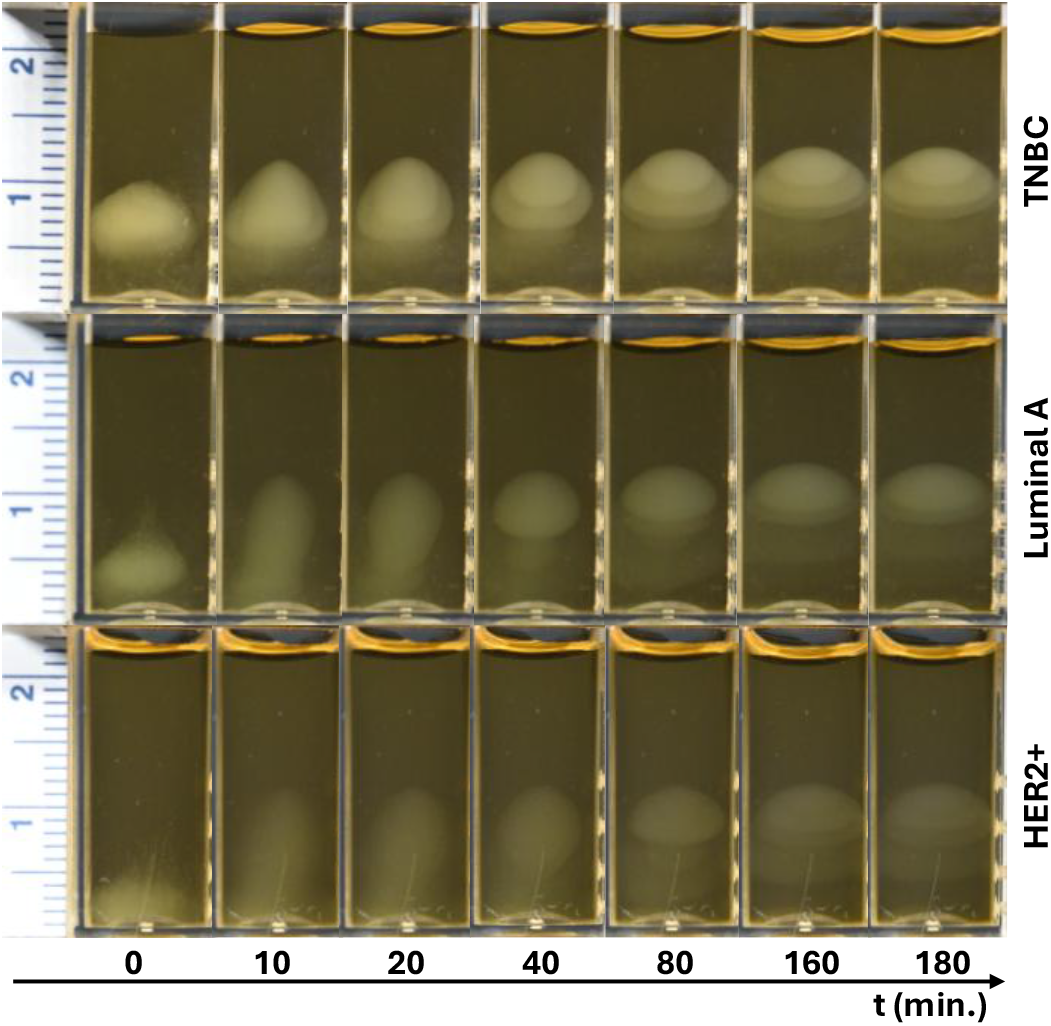
Time-dependent representative levitation patterns of plasma samples from three BC subtypes indicating subtype dependent plasma patterns.

Untargeted metabolomics analysis (**Figure 4**) was performed on the extracted MagLev plasma layers to determine whether the visually distinct levitation patterns contained subtype dependent molecular information. Since cancer associated metabolic reprogramming is known to alter multiple interconnected pathways including glycolysis, amino acid metabolism, lipid remodeling, redox balance, and biosynthetic activity,[13, 53] we hypothesized that the spatially resolved plasma bands generated within the MagLev system may capture subtype specific metabolic signatures beyond conventional single biomarker approaches.[12] Therefore, by combining physical plasma organization within externally applied magnetic field gradients with downstream metabolomics analysis, we aimed to determine whether distinct “metabolomic fingerprints” associated with different BC subtypes could be identified from the levitated plasma patterns.

**Figure 4:**
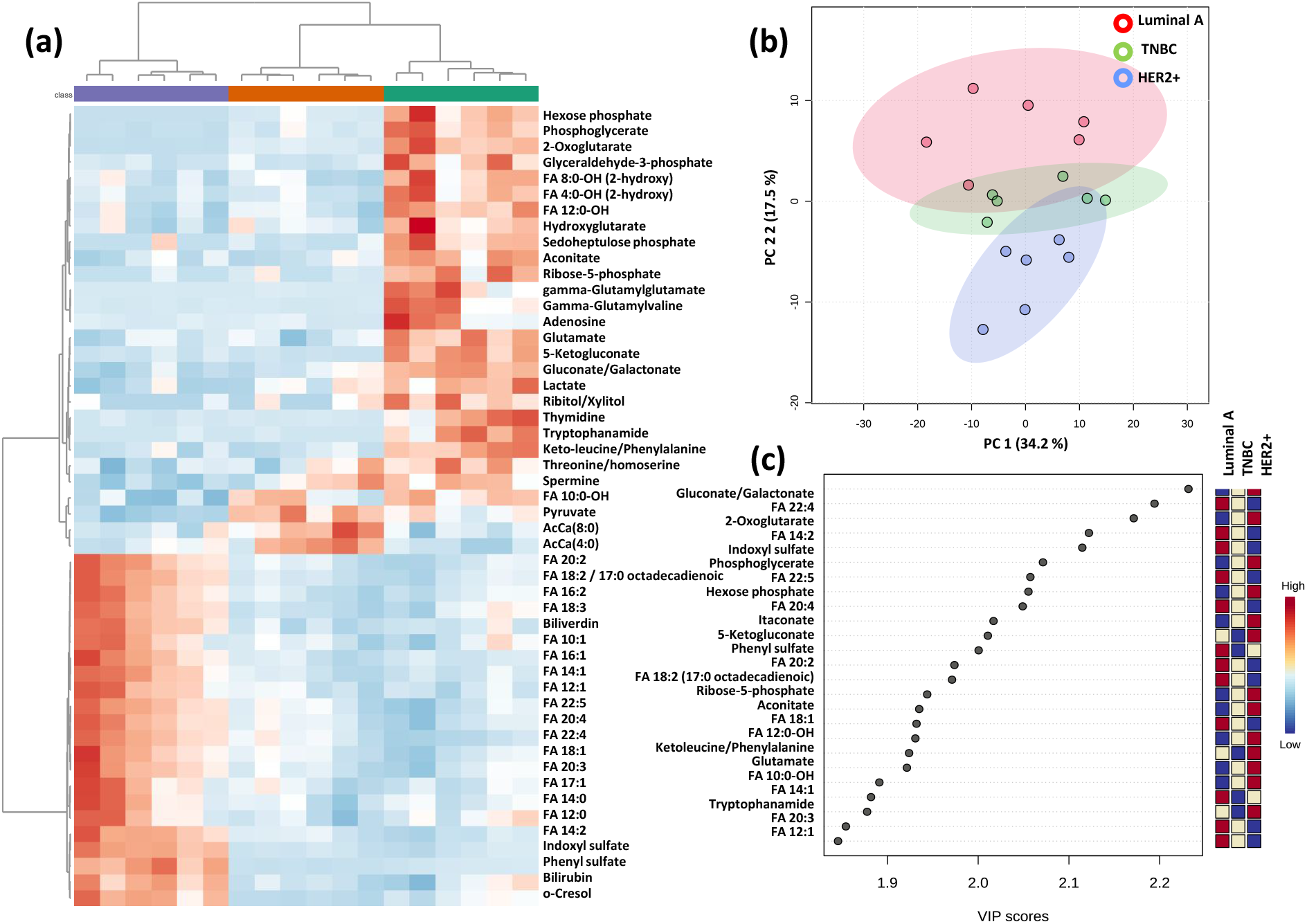
Untargeted metabolomics analysis of levitated plasma patterns from Luminal A, TNBC, and HER2+ BC subtypes. (a) Heatmap analysis of the top 50 altered metabolites reveals distinct subtype dependent metabolomic clustering patterns and coordinated differences in metabolite abundance profiles across the three BC subtypes, (b) PCA demonstrates subtype associated clustering trends and global metabolomic differences among Luminal A, TNBC, and HER2+ samples, and (c) VIP analysis identifies the top metabolites contributing most strongly to BC subtype discrimination, including metabolites associated with glycolysis, TCA cycle activity, amino acid metabolism, redox balance, and lipid metabolism.

The heatmap analysis (**Figure 4a**) provided the first global view of metabolite abundance differences across TNBC, HER2 positive, and Luminal A samples. Clustering of the top altered metabolites showed clear subtype associated patterns, indicating that the levitated plasma layers preserve measurable biochemical differences among the three breast cancer subtypes. The heatmap also showed that these differences were not restricted to a single metabolite or a small isolated group of features. Instead, coordinated changes were observed across several metabolite classes, including central carbon metabolites, amino acid related metabolites, sulfated/aromatic metabolites, and fatty acid species. This suggests that MagLev derived plasma patterning captures broader subtype specific metabolic fingerprints rather than individual biomarkers alone. See **supplementary Table 1** for complete detected metabolite list and normalized metabolite intensity matrix across all samples. PCA was then used as an unsupervised method to evaluate whether the global metabolomics profiles showed natural grouping without using subtype labels during model construction. The PCA plot (**Figure 4b**) showed subtype dependent clustering trends, although partial overlap was observed among some groups. This overlap is not unexpected in a small exploratory dataset and likely reflects both biological heterogeneity among breast cancer subtypes and the unsupervised nature of PCA. See **supplementary Table 2** for significantly altered metabolites identified by ANOVA analysis. This is while PLS DA (**Figure 5a**) provided clearer separation among TNBC, HER2 positive, and Luminal A samples, indicating that the metabolomics profiles contain sufficient subtype related information for supervised discrimination. Unlike PCA, PLS DA uses group information to identify latent variables that maximize separation among predefined classes. It is noteworthy that, although PLS DA separation indicates the feasibility of classifying BC subtypes using MagLev derived metabolomic fingerprints, larger cohorts will be needed for validation. Expanded heatmap analysis of the top 100 altered metabolites (**Figure S2** of supplementary Information (SI)) further demonstrated widespread subtype dependent metabolic reprogramming across multiple interconnected pathways including glycolysis, amino acid metabolism, TCA cycle activity, redox associated metabolites, and lipid remodeling. The broader clustering patterns suggest that BC subtype discrimination is driven by coordinated metabolic network level alterations rather than isolated metabolite changes. Pairwise heatmap comparisons between individual BC subtypes (**Figures S2–S4**) further confirmed distinct metabolomic differences and subtype specific clustering behavior across the levitated plasma patterns.

**Figure 5:**
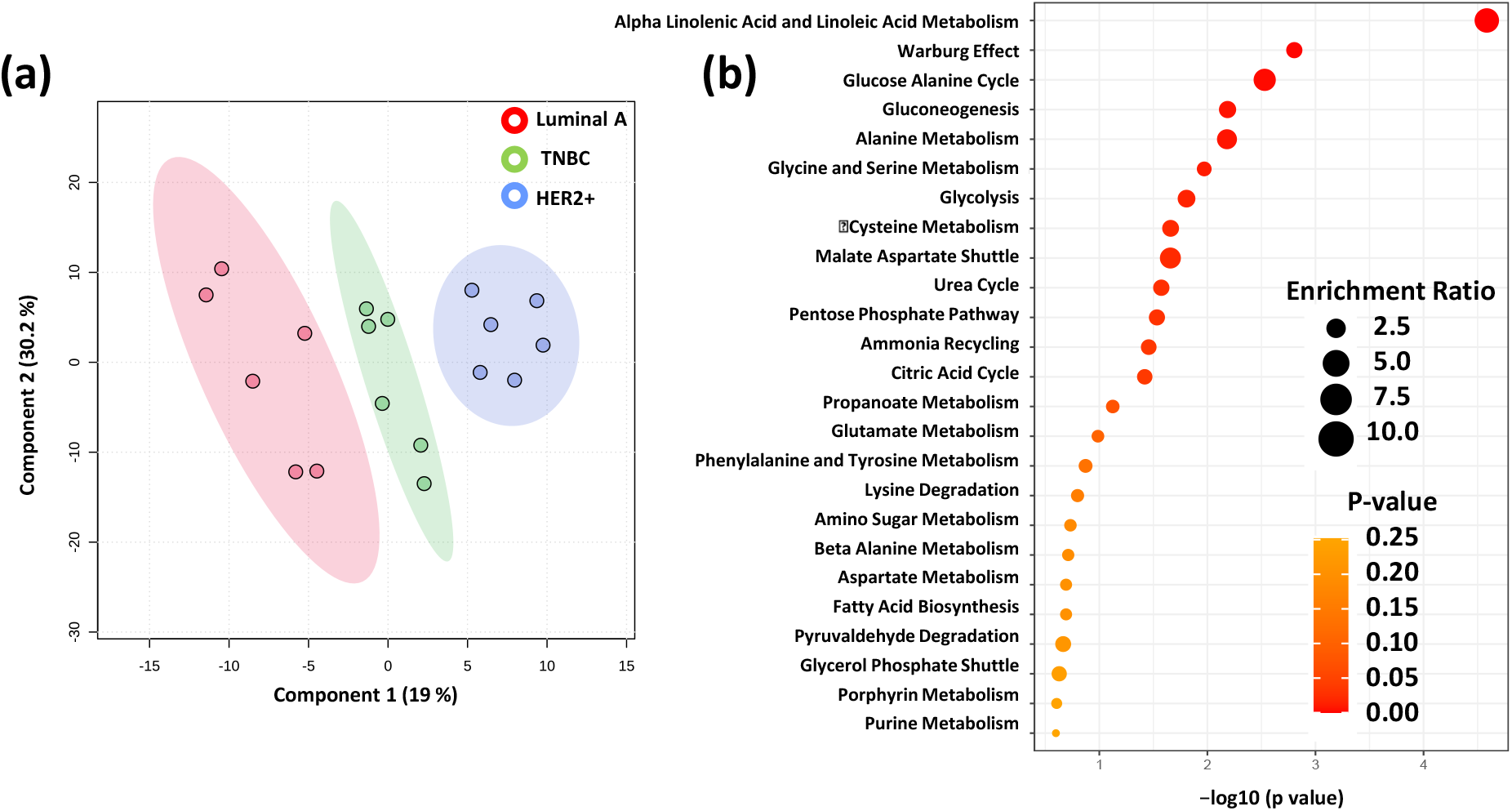
Supervised metabolomics analysis and pathway enrichment of levitated plasma patterns from Luminal A, TNBC, and HER2+ BC. (a) PLS DA analysis demonstrates clear subtype dependent separation among Luminal A, TNBC, and HER2+ samples based on global metabolomic profiles obtained from the levitated plasma layers. The supervised clustering indicates that the extracted plasma patterns contain sufficient metabolomic information to discriminate among biologically distinct BC subtypes with minimal overlap between groups, and (b) metabolite pathway enrichment analysis of the top altered metabolites reveals significant subtype associated metabolic reprogramming across multiple interconnected pathways, including the Warburg effect, glycolysis, gluconeogenesis, pentose phosphate pathway, citric acid cycle, amino acid metabolism, glutamate metabolism, fatty acid biosynthesis, and redox associated pathways.

VIP analysis (**Figure 4c**) identified the metabolites contributing most significantly to the PLS DA based subtype separation. The top discriminating metabolites included gluconate or galactonate, FA 22:4, 2 oxoglutarate, FA 14:2, indoxyl sulfate, phosphoglycerate, FA 22:5, hexose phosphate, FA 20:4, itaconate, 5 ketogluconate, phenyl sulfate, ribose 5 phosphate, aconitate, and glutamate. These metabolites suggest that subtype separation is driven by several biologically meaningful pathways, including glycolysis, the pentose phosphate pathway, TCA cycle activity, amino acid metabolism, aromatic sulfate metabolism, and lipid remodeling.[54, 55] This is critical as it shows that the observed separation of magnetically BC plasma samples is not only statistical, but also biologically interpretable.

The pathway enrichment analysis (**Figure 5b**) further supports this interpretation. Enriched pathways included glycolysis, gluconeogenesis, pentose phosphate pathway metabolism, citric acid cycle related metabolism, glutamate metabolism, and fatty acid metabolism. These pathways are closely linked to cancer cell proliferation, redox balance, biosynthesis, and membrane lipid remodeling.[55] HER2 positive samples showed stronger association with energy metabolism related features, including metabolites connected to glycolysis, pentose phosphate pathway activity, and TCA cycle metabolism.[56] Luminal A samples showed stronger lipid associated signatures, especially long chain fatty acid related features, suggesting subtype dependent lipid remodeling.[57] TNBC showed a distinct but partially intermediate metabolic profile, which is consistent with its biological heterogeneity and aggressive phenotype.[4]

To further validate the subtype dependent metabolic differences, we next performed pairwise volcano plot analysis (**Figures S5–S7**) between individual BC subtype comparisons. Although the previous analyses provided an overall systems level view of subtype associated metabolic reprogramming, the volcano plots enabled visualization of specific metabolites showing the strongest fold change and statistical significance between individual subtype pairs. These analyses further confirmed distinct metabolomic differences among Luminal A, TNBC, and HER2+ samples and identified subtype specific metabolites contributing to the observed metabolic heterogeneity. Several metabolites demonstrated statistically significant fold changes between individual subtype comparisons, supporting distinct metabolic reprogramming among the three BC subtypes. Differentially abundant metabolites included features associated with glycolysis, TCA cycle intermediates, amino acid metabolism, aromatic metabolites, and lipid metabolism, further supporting the metabolomic heterogeneity observed in the global clustering and pathway enrichment analyses.[56] These findings indicate that BC subtype discrimination is driven by coordinated metabolic reprogramming involving energy metabolism, glycolysis, amino acid metabolism, redox balance, and lipid remodeling rather than isolated metabolite changes alone.[55] Importantly, the results demonstrate that the distinct and dynamically evolving MagLev plasma patterns retain biologically meaningful molecular information associated with different breast cancer subtypes. While conventional bulk plasma analysis often averages molecular complexity into a single metabolic profile, the MagLev platform enables spatial organization and fractionation of plasma biomolecules into distinct levitated layers with subtype dependent metabolomic signatures. This suggests that disease associated metabolic heterogeneity can be partially resolved through magnetic field driven plasma organization prior to downstream omics analysis. These findings support the potential of combining MagLev based plasma patterning with untargeted metabolomics analysis as a promising strategy for generating subtype specific metabolomic fingerprints for breast cancer discrimination.[58]

## Conclusions

In summary, this exploratory proof-of-concept study demonstrates that magnetic levitation combined with untargeted metabolomics analysis can capture subtype dependent molecular differences among Luminal A, TNBC, and HER2+ BC plasma samples. Distinct and dynamically evolving MagLev plasma patterns, together with global metabolomics profiling, revealed reproducible subtype associated metabolic signatures involving glycolysis, amino acid metabolism, TCA cycle activity, redox balance, and lipid remodeling. Although the current study is limited by the small sample size and requires validation in larger patient cohorts, the findings establish the feasibility of MagLev derived metabolomic fingerprinting as a potential platform for BC subtype discrimination. Future studies involving expanded clinical cohorts, longitudinal sampling, and integrated multi omics analysis may further improve the robustness, biological interpretation, and translational potential of this approach for disease classification and biomarker discovery.

## Associated content

### Supporting Information

Supporting figures/tables and LC-MS/MS analyses.

## Authors’ information

### Notes

The authors declare no competing financial interest.

## Acknowledgments

S.V and A.A.A acknowledge financial support from National Institutes of Health (grant number R03EB034817) and UCCS CRCW seed grant.

