## Supporting Information for "Magnetic Levitation Derived Metabolomic Fingerprinting Enables Exploratory Discrimination of Breast Cancer Subtypes"

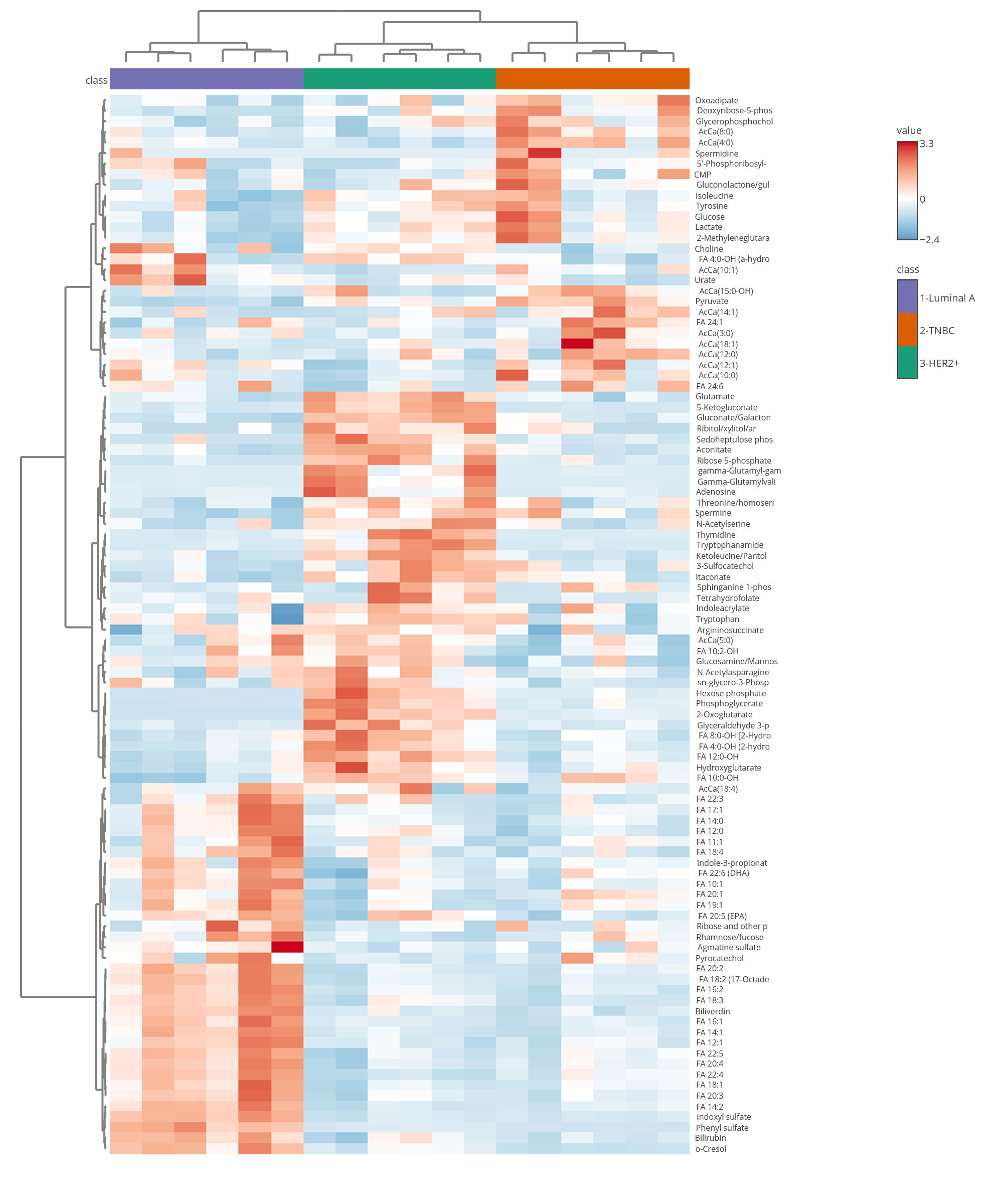

**Figure S1:** Expanded heatmap analysis of the top 100 altered metabolites identified from untargeted metabolomics analysis of levitated plasma patterns from Luminal A, TNBC, and HER2+ breast cancer subtypes. Hierarchical clustering reveals broad subtype dependent metabolomic differences and coordinated changes across multiple interconnected metabolic pathways, including glycolysis, pentose phosphate pathway activity, amino acid metabolism, TCA cycle intermediates, redox associated metabolites, aromatic metabolites, and lipid remodeling. The larger metabolite set further supports the presence of distinct subtype specific metabolomic fingerprints and demonstrates that BC subtype discrimination arises from global metabolic network alterations rather than isolated metabolite changes alone.

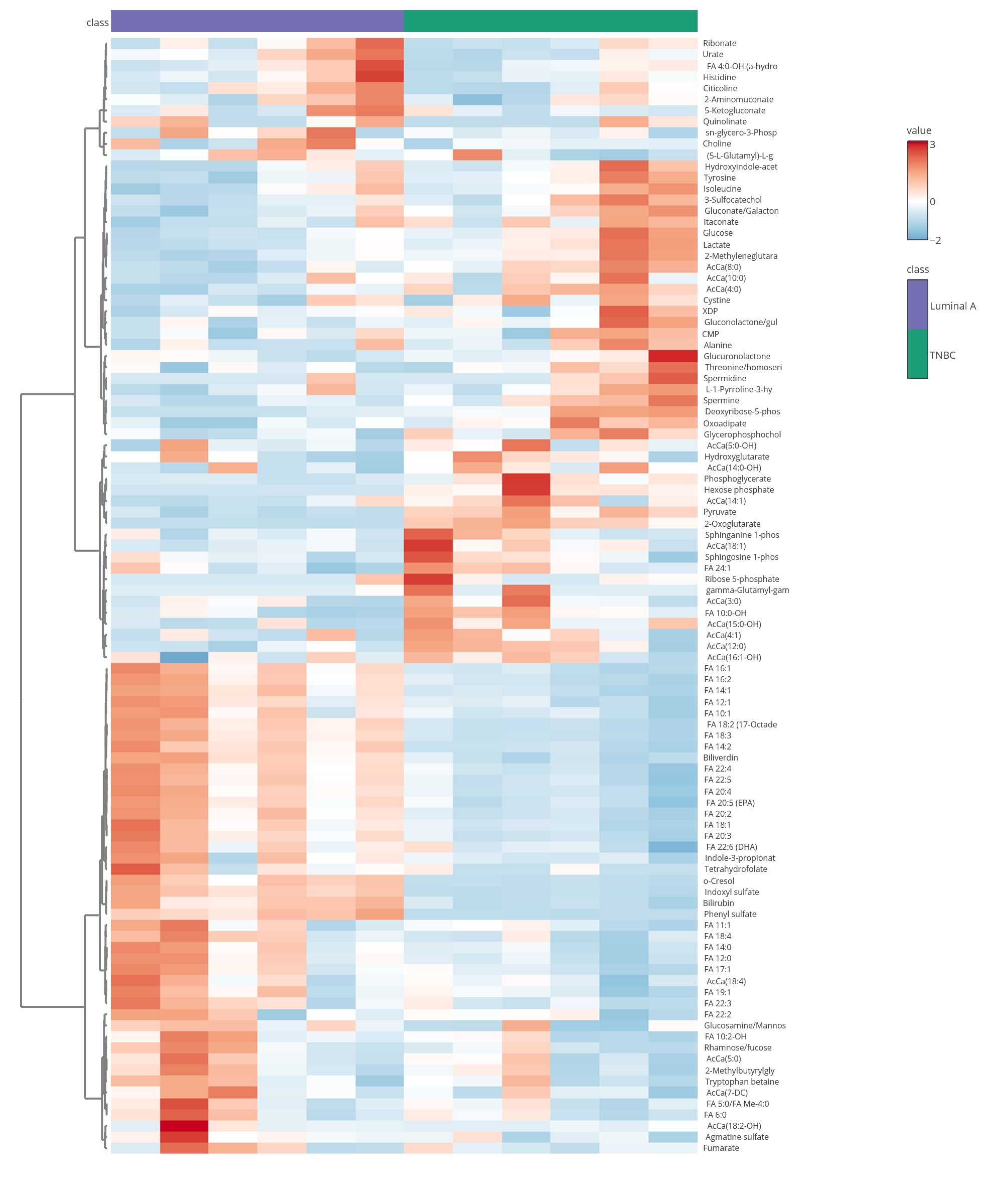

**Figure S2:** Pairwise heatmap analysis of the top altered metabolites between Luminal A and TNBC BC subtypes.

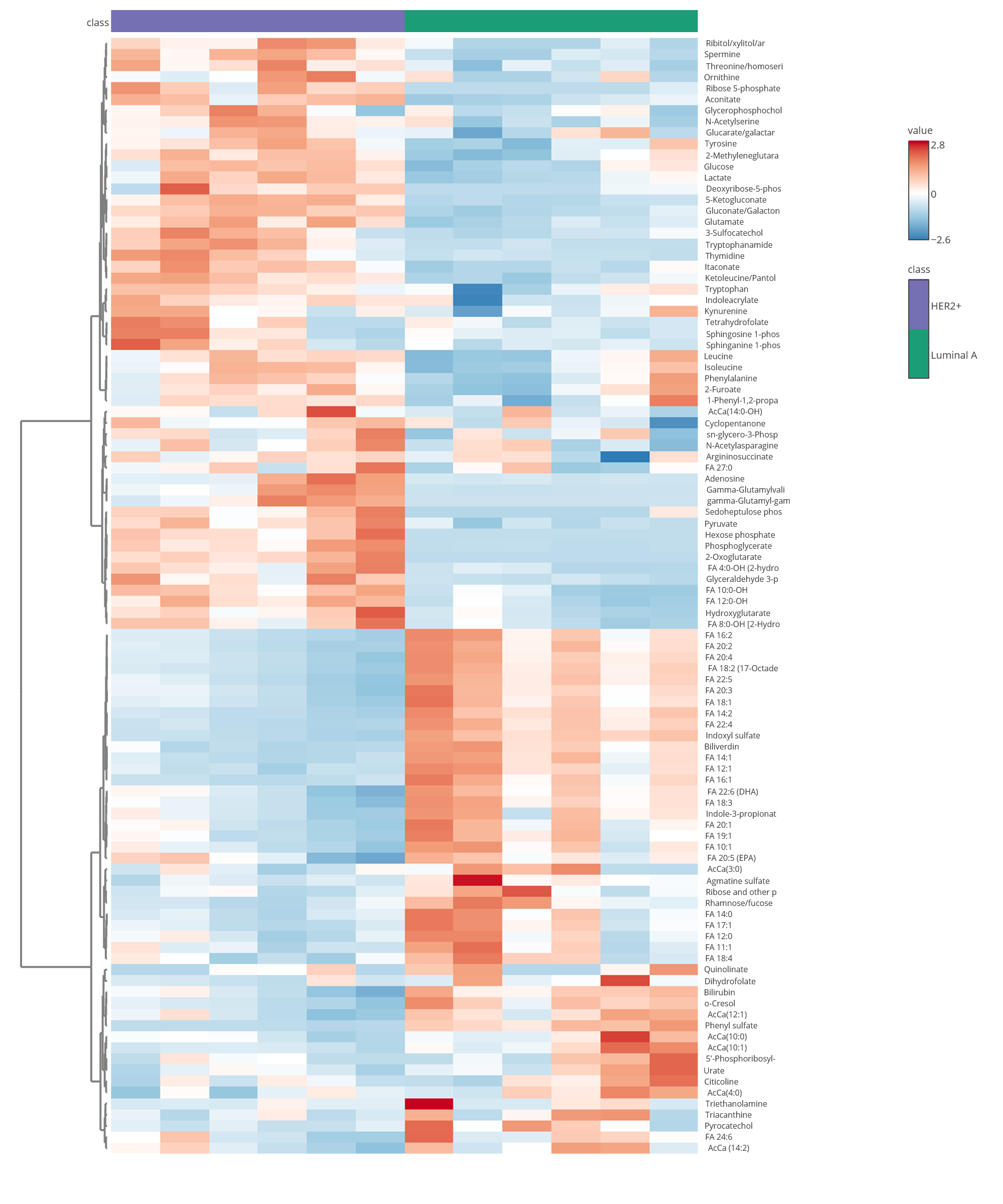

**Figure S3:** Pairwise heatmap analysis of the top altered metabolites between Luminal A and HER2+ BC subtypes.

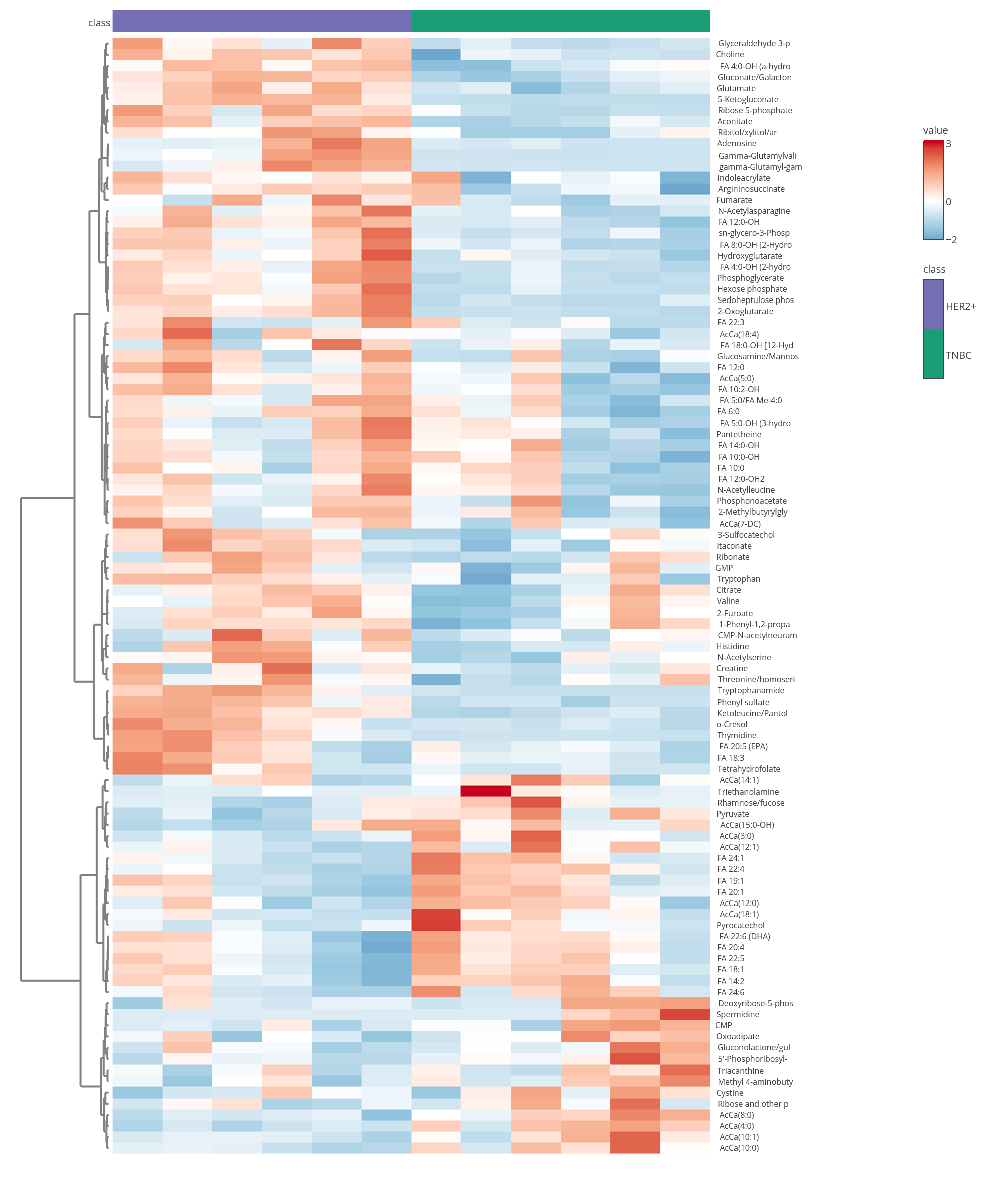

**Figure S4:** Pairwise heatmap analysis of the top altered metabolites between HER2+ and TNBC BC subtypes.

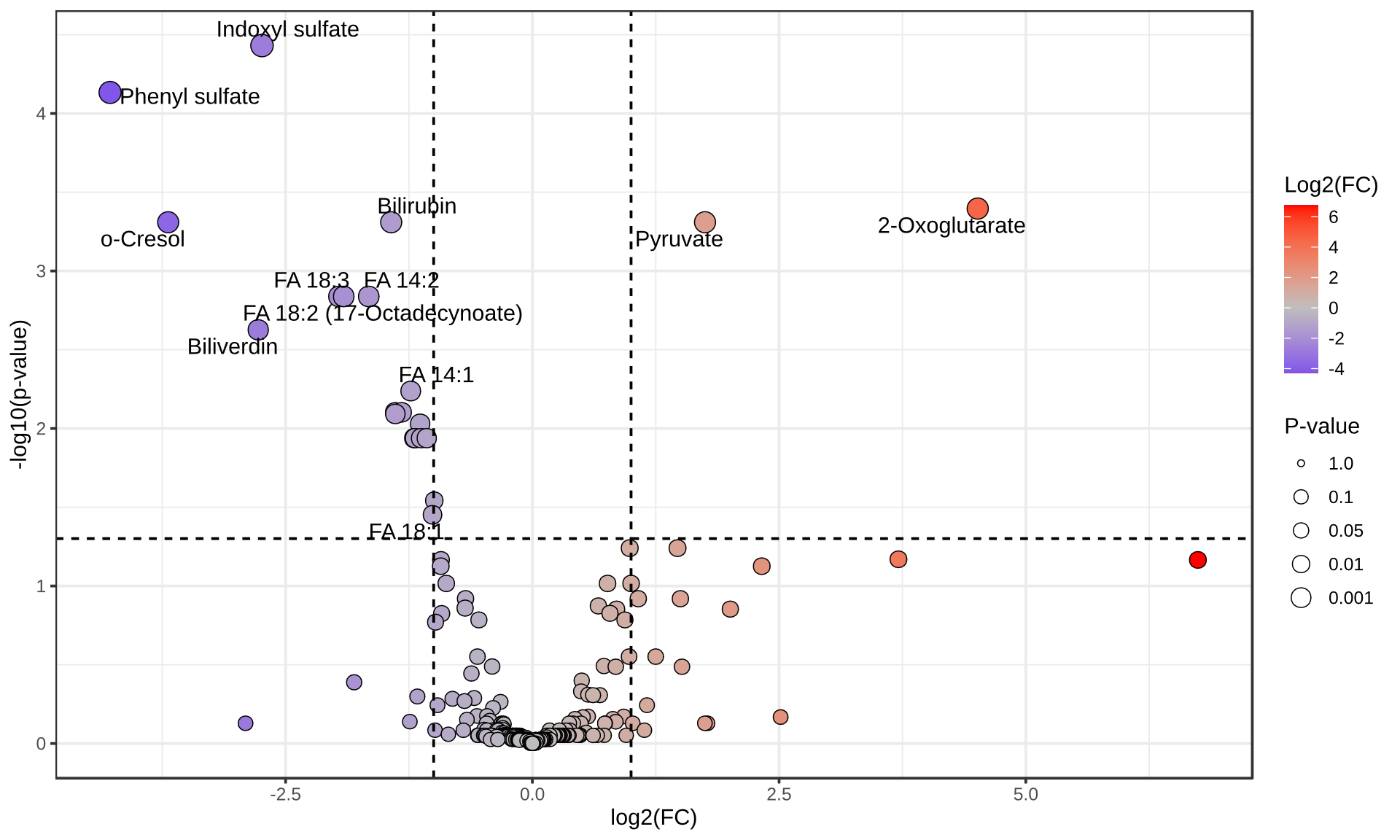

**Figure S5:** Pairwise volcano plot analysis comparing Luminal A and TNBC BC subtypes. Significantly altered metabolites showing differential fold changes and statistical significance between the two subtypes are highlighted, further supporting subtype dependent metabolomic differences within the levitated plasma patterns.

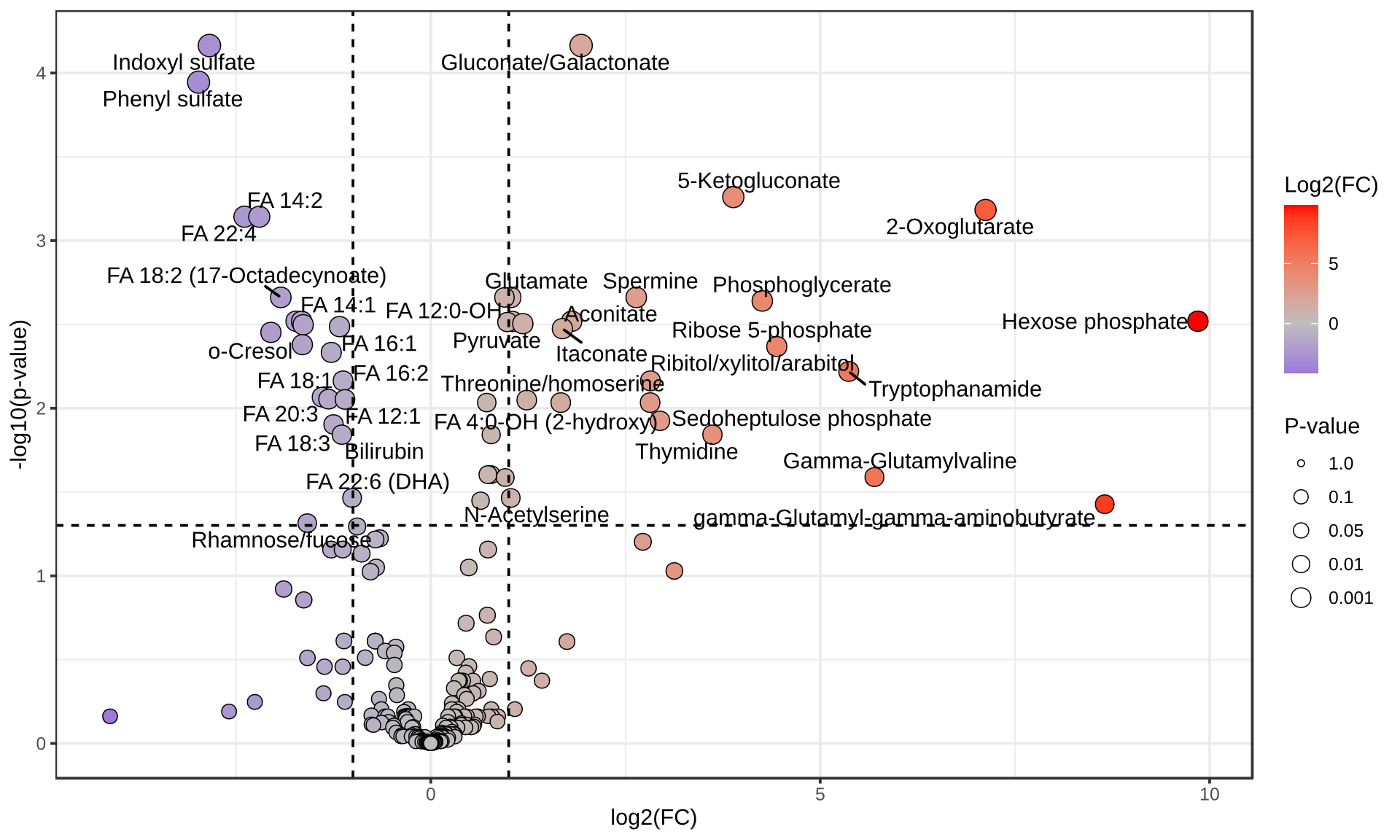

**Figure S6:** Pairwise volcano plot analysis comparing Luminal A and HER2+ BC subtypes.

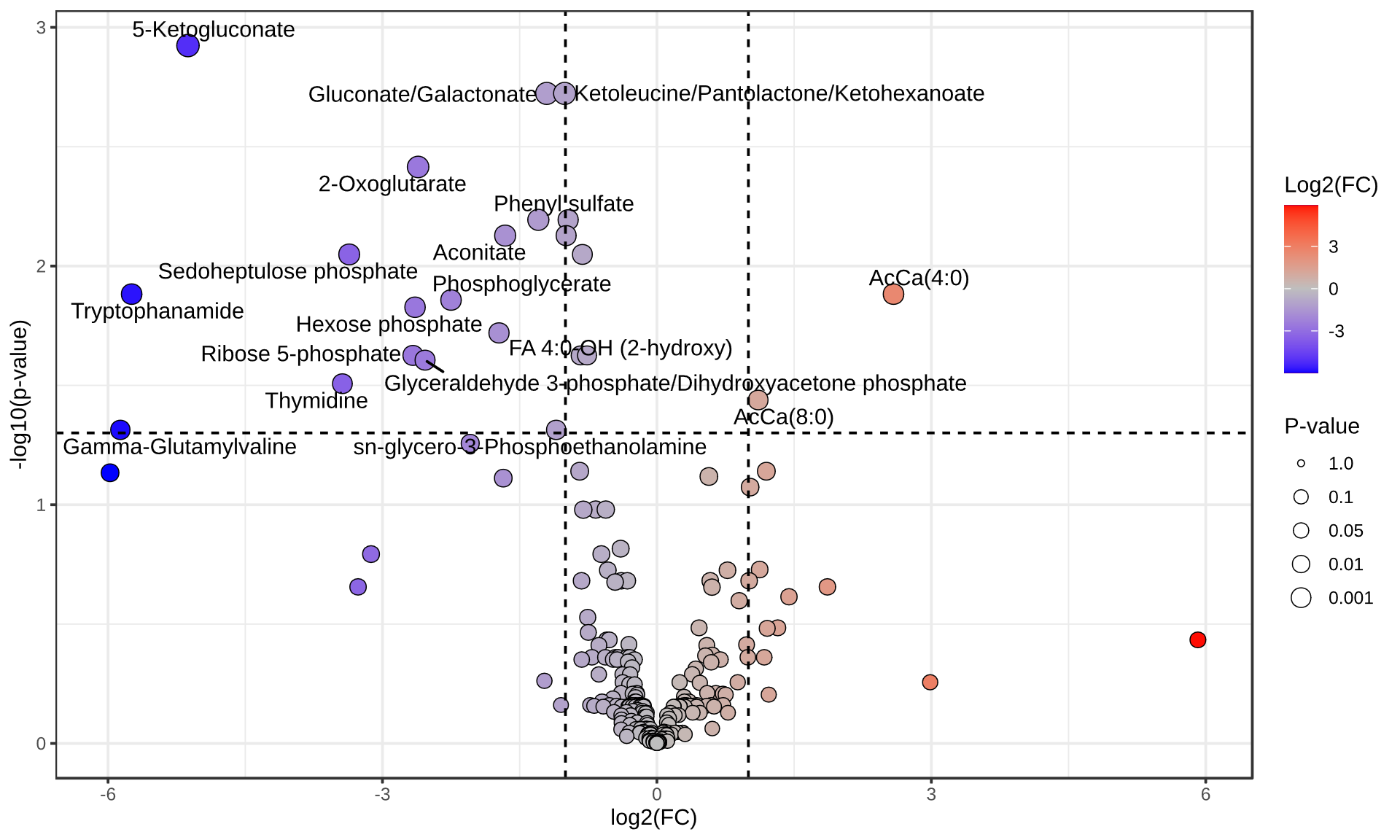

**Figure S7:** Pairwise volcano plot analysis comparing TNBC and HER2+ BC subtypes.
